# Evolving patterns of extreme publishing behavior across science

**DOI:** 10.1101/2023.11.23.568476

**Authors:** John P.A. Ioannidis, Thomas A. Collins, Jeroen Baas

## Abstract

Extreme publishing behavior may reflect a combination of some authors with genuinely high publication output and of other people who have their names listed too frequently in publications because of consortium agreements, gift authorship or other spurious practices. We aimed to evaluate the evolution of extreme publishing behavior across countries and scientific fields during 2000-2022. Extreme publishing behavior was defined as having >60 full articles (original articles, reviews, conference papers) in a single calendar year and indexed in Scopus. We identified 3,191 authors with extreme publishing behavior across science excluding Physics and 12,624 such authors in Physics. While Physics had much higher numbers of extreme publishing authors in the past, in 2022 extreme publishing authors was almost as numerous in non-Physics and Physics disciplines (1,226 vs. 1,480). Excluding Physics, China had the largest number of extreme publishing authors, followed by the USA. The largest fold-wise increases between 2016 and 2022 (5-19-fold) occurred in Thailand, Saudi Arabia, Spain, India, Italy, Russia, Pakistan, and South Korea. Excluding Physics, most extreme publishing authors were in Clinical Medicine, but from 2016 to 2022 the largest relative increases (>6-fold) were seen in Agriculture, Fisheries & Forestry, Biology, and Mathematics and Statistics. Extreme publishing authors accounted for 4,360 of the 10,000 most-cited authors (based on raw citation count) across science. While most Physics authors with extreme publishing behavior had modest citation impact in a composite citation indicator that adjusts for co-authorship and author positions, 67% of authors with extreme publishing behavior in non-Physics fields remained within the top-2% according to that indicator among all authors with >=5 full articles. Extreme publishing behavior has become worryingly common across scientific fields with rapidly increasing rates in some countries and settings and may herald a rapid depreciation of authorship standards.

## INTRODUCTION

Authorship of scientific articles is highly coveted in the “publish or perish” mentality. Many scientists are very active, publishing large numbers of articles each year. Having one’s name listed in huge numbers of publications is facilitated further by the advent of team science, especially in fields where extremely large numbers of authors are listed in each article, and by an inflation of authors due to changes in norms of credit allocation (Hosseini et al., 2022; Wager et al., 2015; Papatheodorou et al., 2008). Extreme publishing behavior means being frequently listed as an author, regardless of whether authorship criteria are formally met. In this article we define extreme publishing behavior using empirical data and coining a standardized threshold of very high annual productivity within any calendar year. Authors with a publication record spanning many decades may accumulate a large number of authored articles and this would not be very surprising. Conversely, concentrations of very high number of published articles within a single calendar year is more likely to raise questions about the nature and legitimacy of authorship contributions for massive work that is so close time-wise.

Extreme publishing behavior may reflect a combination of genuinely extremely capable (at publishing) authors and others who may have their names listed in very large numbers of articles because of consortium agreements, gift authorship (e.g. to department heads and lab directors) or other spurious patterns. For people who demonstrate extreme publishing behavior, it is often difficult to separate to what extent their records may reflect true capability to generate large publication output with genuine author contributions, lenient criteria for authorship credit or spurious research and authorship practices and even fraud. Studying the most extreme cases may help understand the evolving dynamics of research publishing and authorship.

It is arbitrary to set a threshold of what is extreme. However, previous work (Ioannidis et al., 2018) defined as “hyperprolific” authors as those who, in any single calendar year, had published more than 72 full articles (including original articles, reviews, and conference papers and excluding editorials/commentaries, notes, and letters). Such publishing records amounts to publishing more than 1 full article every 5 days, even counting weekends. Analysis of the Scopus data for the period 2000-2016 (Ioannidis et al, 2018) found that the vast majority of these authors were in physics disciplines, nuclear and particle physics specifically, reflecting the well-known practice in that field of equating authorship with participation in some experimental group without necessarily writing or editing the resulting articles. Excluding physics, during 2000-2016 the few hyperprolific authors (n=154, excluding Chinese names) were mostly concentrated in a few scientific subfields such as epidemiology, cardiology, and crystallography (Ioannidis et al, 2018). Their numbers had clearly increased between 2000 and 2014, but seemed to level off between 2014 and 2016. An interview survey of these authors revealed that frequently there was a lax approach towards traditional Vancouver authorship criteria (Ioannidis et al, 2018).

Since 2016, the pressure to publish or perish may have grown stronger across several scientific fields. Extreme publishing behavior has also been further incentivized in some countries and settings, including monetary benefits which are sometimes out of proportion to typical salaries of researchers (Xu et al., 2021; Quan et al., 2017; 7. Kim & Bak, 2016; Andersen & Pallesen, 2008; Chapman et al., 2019). Some scientists with extreme publishing behavior even adopt spurious affiliations with Saudi Arabian universities to secure financial benefits (Bhattacharjee, 2011; Catanzaro, 2023). Concurrently, the paradigm of team science has become more common across an increasing number of scientific fields (Fontanarosa et al., 2017). The effect these evolving circumstances have had on the phenomenon of extreme publishing behavior is unknown. It would be also be interesting to examine which countries and scientific fields are particularly affected. Therefore, in this article we examined the evolving presence of extreme publishing behavior across science in the extended period 2000-2022.

The rest of the article is organized as follows. First, we present the Methods, including the databases used and definitions; and the analyses that we specified. Then, we present the respective Results of these analyses. Finally, Conclusions are offered including placing the current work in context of other evidence, discussing some limitations, and addressing implications for the evolution of publishing and authorship and future work.

## METHODS

The Methods section present the key features of the databases that we used, and the definitions that we set for identifying an author as having extreme publishing behavior in a given calendar year. Then, the analyses are specified, including the evolution of extreme publishing behavior over time in 2000-2022, as well as analyses at the level of each country and at the level of each scientific field.

### Databases and definitions

We used the Scopus database (Baas, et al. 2020) with a data freeze on May 2023. The full period 2000-2022 (23 calendar years) was considered. Similar to previous work (Ioannidis et al, 2018), we considered the number of full articles published by each scientist. Full articles included the Scopus categories “article” (original articles), “review”, and “conference paper”. All other items were not counted (including the categories “editorial”, “note”, “letter”, “correction”, and others). Of note, the number of hyperprolific authors identified in the previous work for the years 2000-2016 is expected to be different in the current analysis, because more journals have been added to Scopus retrospectively for these years, some published items may have changed characterization, and author IDs are continuously corrected for errors (e.g. some authors who had their articles split in two or more separate Scopus ID files may have had their files combined and thus now they would emerge as fulfilling extreme publishing behavior criteria, while this was not previously apparent). Eligible authors were selected based on the number of full articles that they had published in a single calendar year exceeding a set threshold.

As previously defined (Ioannidis et al., 2018), an author was called “hyperprolific” for a given calendar year if he/she had their names listed as an author in more than 1 full article every 5 days, i.e. 73 or more full articles. Moreover, we extended the capture of authors who have their names appear in many articles to also take into account authors who have published 61-72 full articles in any given year, i.e. more than 1 article every 6 days (>60) but not more than 1 article every 5 days (<73) (hence called “almost hyperprolific”. The sum of “hyperprolific” and “almost hyperprolific” authors are called extreme publishing authors (authors with extreme publishing behavior). This lower threshold (>60) may allow capturing a broader set of authors who probably follow overtly lenient, spurious or even fraudulent publication and authorship practices, although it is possible that some of even many of them are simply genuinely extremely productive.

For each Scopus author ID that met the criteria for extreme publishing behavior in a calendar year, we captured the number of full articles in that calendar year, the Scopus listed affiliation and country of affiliation (at the time of the Scopus data freeze), the total number of full articles published in his/her career and during 2000-2022, and the main scientific subfield of his/her work.

Scientific fields were classified according to the Science-Metrix classification (Archambault et al., 2011) into 20 fields and 174 subfields. The classification is principally journal-based, with each journal allocated to one field/subfield, except for the minority of multidisciplinary journals where published items may be allocated to more than one field/subfield. For each author, the main field is the one where he/she has published more items during 2000-2022. In the case where the author contributed equal output to two or more fields, the subfield in which the author had the highest amount of publications relative to the total number of publications in that field is chosen as the author’s field.

We cross-linked these data with the data from another project (Ioannidis et al., 2019) where we have generated for all authors with at least 5 full articles information on 6 citation indicators (total citations, h-index, co-authorship-adjusted hm index, citations to single authored items, citations to single or first authored items, citations to single/fist/last authored items) and a composite citation indicator combining these 6 indicators (Ioannidis et al., 2016). Authors are then ranked based on the composite citation indicator across all authors and also specifically across all the authors allocated to the same main scientific subfield. Percentile ranking is then provided for each author across science and within his/her main subfield.

The composite indicator adjusts for co-authorship and author positions and is thus expected to attenuate the relative impact and ranking of many authors with extreme publishing behavior, especially in middle-author positions. Composite indicator values and rankings are typically updated by our team every year (Ioannidis et al., 2019); however, for this project we also separately updated them specifically using the May 2023 Scopus data freeze that was used also for the collection of hyperprolific almost hyperprolific and extreme publishing data, so that the linked data would be consistent.

### Analyses

We began by separating upfront authors whose main subfield is one of the subfields of the Physics & Astronomy field in the Science-Metrix classification (henceforward called “Physics” group for parsimony), from those whose main subfield is within one of the other 19 fields of science (“non-Physics” group). This was essential, since it is well documented that the large majority of extreme publishing authors have traditionally been in physics-related disciplines (Ioannidis et al., 2018). The main analyses focused on the non-Physics group, but we also examined the evolution of extreme publishing behavior in the Physics group for contrast.

The main analyses examined the number of hyperprolific, almost hyperprolific, and extreme publishing authors in each calendar year between 2000-2022 to discern and describe time patterns. We also evaluated the distribution of these authors in different countries and assessed whether the rate of increase in such authors is particularly high in recent years (2016-2022) in specific countries. Furthermore, we evaluated the distribution of these authors each year in the main fields of science. This allowed us to discern and describe whether the rate of increase in such authors is particularly high in recent years (2016-2022) in specific fields. We then explored whether any specific subfield(s) within the fields with most rapidly increasing presence of such authors is responsible for the massive increase. Rates of authors with extreme publishing behavior were also expressed in conjunction with the total number of authors with >=5 full articles during 2000-2022 in each country and in each field.

In order to describe the citation impact of hyperprolific, almost hyperprolific, and extreme publishing authors, we evaluated how many of them were ranked among the top-10,000 most-cited scientists across non-Physics and Physics based on raw citation counts including self-citations; and in the top-2% percentile according to the composite citation indicator within their main subfield (with or without self-citations). We also generated boxplots of the percentile rankings within their subfield for such authors, so as to visualize the features of the full distribution of composite citation indicator-based rankings. These analyses allow to understand whether authors with extreme publishing behavior comprise a large share of those who are ranked among those with highest citation impact.

For the Physics group, we also examined whether changes over time in the total number of hyperprolific, almost hyperprolific, and extreme publishing authors are related to the number of full articles published every year that include an affiliation from the European Organization for Nuclear Research (EONR) (AFID(60019778)) and even more specifically to those articles among them that have a large number of listed authors (>100, >500, and >1000).

## RESULTS

### Number of scientists with extreme publishing behavior

Table 1 shows for each calendar year in each of these two groups, the number of hyperprolific (>72 full articles published in a single year), almost hyperprolific (61-72 full articles published in a single year), and extreme publishing authors (>60 full articles published in a single year). As defined, the number of extreme publishing authors is the sum of the numbers of hyperprolific and almost hyperprolific authors. During 2000-2022, in the non-Physics group (all scientific fields, excluding the “Physics & Astronomy” field in the Science-Metrix classification), there were 1,661 scientists who had reached hyperprolific status in at least one calendar year, 2,543 authors who had reached almost hyperprolific status in at least one calendar year, and overall 3,191 scientists who had reached extreme publishing status in at least one calendar year. The respective numbers for the Physics group were 10,441, 8,588, and 12,624.

**Table 1.** Number of hyperprolific, almost hyperprolific and overall extreme publishing authors in 2000-2022 in Physics and non-Physics groups.

| Year | Hyperprolific<br>Physics | Hyperprolific<br>non-Physics | Almost<br>hyperprolific<br>Physics | Almost<br>hyperprolific<br>non-Physics | Eextreme<br>publishing<br>Physics | Extreme<br>publishing<br>non-Physics |
| --- | --- | --- | --- | --- | --- | --- |
| 2000 | 7 | 19 | 9 | 16 | 16 | 35 |
| 2001 | 5 | 10 | 8 | 29 | 13 | 39 |
| 2002 | 12 | 13 | 22 | 25 | 34 | 38 |
| 2003 | 15 | 24 | 17 | 34 | 32 | 58 |
| 2004 | 25 | 49 | 84 | 53 | 109 | 102 |
| 2005 | 36 | 64 | 117 | 74 | 153 | 138 |
| 2006 | 114 | 93 | 500 | 78 | 614 | 171 |
| 2007 | 501 | 108 | 124 | 79 | 625 | 187 |
| 2008 | 50 | 105 | 473 | 110 | 523 | 215 |
| 2009 | 44 | 132 | 217 | 106 | 261 | 238 |
| 2010 | 125 | 126 | 187 | 130 | 312 | 256 |
| 2011 | 958 | 143 | 1513 | 161 | 2471 | 304 |
| 2012 | 5170 | 141 | 215 | 149 | 5385 | 290 |
| 2013 | 4985 | 137 | 340 | 162 | 5325 | 299 |
| 2014 | 4580 | 169 | 826 | 204 | 5406 | 373 |
| 2015 | 4669 | 177 | 771 | 200 | 5440 | 377 |
| 2016 | 4938 | 185 | 252 | 202 | 5190 | 387 |
| 2017 | 4863 | 206 | 770 | 235 | 5633 | 441 |
| 2018 | 5039 | 294 | 213 | 313 | 5252 | 607 |
| 2019 | 4935 | 360 | 361 | 355 | 5296 | 715 |
| 2020 | 4155 | 431 | 686 | 414 | 4841 | 845 |
| 2021 | 972 | 568 | 3944 | 572 | 4916 | 1140 |
| 2022 | 761 | 674 | 719 | 592 | 1480 | 1266 |

The Physics group witnessed a very sharp increase in the number of hyperprolific authors between 2010 (n=125) and 2012 (n=5,170) and the number of hyperprolific authors remained relatively constant at 5,000 until 2019. In 2020 there was a small decrease and in 2021 and 2022 there was a very sharp decrease; in 2021, the sharp decrease in hyperpprolific authors was compensated by an equally sharp increase in the number of almost hyperprolific authors, but this compensation did not seem to occur in 2022. This pattern is largely explained by examination of the full articles published each year with the affiliation of European Organization of Nuclear Research. The number of such articles has declined sharply in the last few years, with the greatest decline in 2022 and with even greater proportional declines for the number of articles with this affiliation who have a large number of authors (>100, >500, >1000) (supplementary table 1 and supplementary figure 1)

Excluding Physics, the number of both hyperprolific and almost hyperprolific authors showed a 5-fold increase between 2000 and 2006, it increased 2-3-fold in the next decade and has seen another acceleration of growth with 3-4-fold increase in the last 6 years (2016-2022). In 2022, the number of hyperprolific and almost hyperprolific authors across science excluding Physics was almost similar to the respective numbers in Physics. Excluding Physics, there were 1,266 authors with extreme publishing behavior in 2022 (versus 1,480 for Physics) (Table 1).

### Extreme publishing behavior across science: country-level analyses

Figure 1 shows the share of each country in hyperprolific, almost hyperprolific, and overall extreme publishing authors during the cumulative period 2000-2022 in the non-Physics and Physics groups (detailed numerical data appear in Supplementary Table 2).

**Figure 1.**
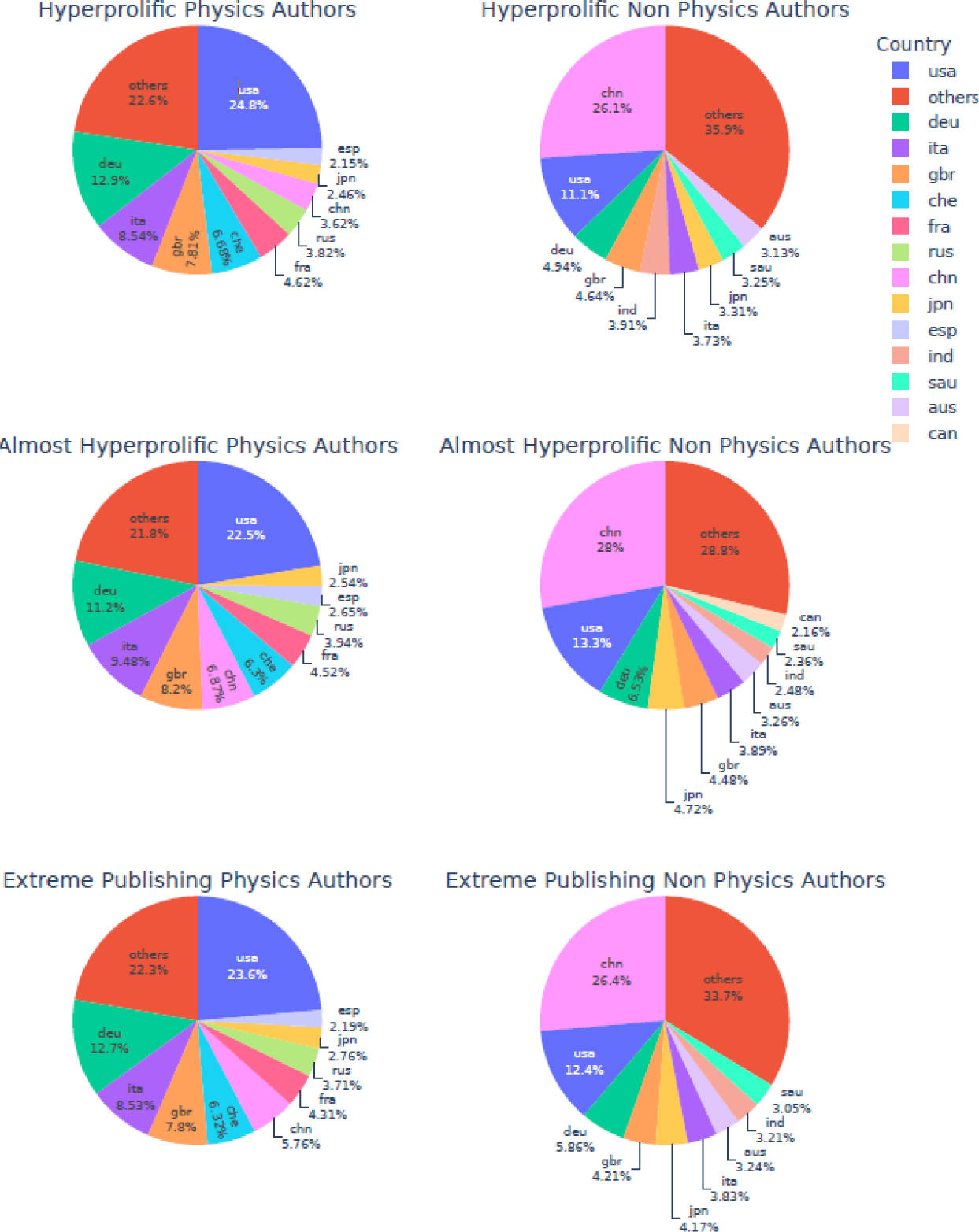
Pie charts of the number of hyperprolific, almost hyperprolific, and extreme publishing (sum of hyperprolific and almost hyperprolific) authors in Physics and non-Physics scientific fields according to country of affiliation for the entire period 2000-2022.

Some countries accounted for the lion’s share of authors with extreme publishing behavior. Patterns are similar for hyperprolific, almost hyperprolific, and extreme publishing authors, but they differ in the non-Physics versus Physics fields. In Physics the main countries reflected to a large extent participation in EONR projects and thus the order was USA, Germany, Italy, UK, Switzerland, China, and France. Excluding Physics, China has had the highest number of extreme publishing authors every year since 2003 and in the cumulative 2000-2022 period it was followed by the USA, Germany, Japan, UK, Italy, and Australia. However, in recent years some countries had risen much faster towards the top ranks in the non-Physics fields.

In non-Physics fields during 2022 specifically, the countries with the highest number of authors with extreme publishing behavior were China (n=303), USA (n=124), Saudi Arabia (n=69), Italy (n=62), Germany (n=58), India (n=51), UK (n=47), Australia (n=47), Japan (n=35), Canada (n=28), Iran (n=26), South Korea (n=26), Spain (n=23), Netherlands (n=20), Taiwan (n=19), Thailand (n=19), Pakistan (n=17), Denmark (n=15), Malaysia (n=14), France (n-14), Russia (n=13), Singapore (n=12), Hong Kong (n=11), and Switzerland (n=7). Compared to 2016, most countries had 1.5-4-fold increases in the numbers of authors with extreme publishing behavior, but several countries had much higher increases: Thailand 19-fold (19 vs. 1), Saudi Arabia 11.5-fold (69 vs. 6), Spain 11.5-fold (23 vs. 2), India 10.2-fold (51 vs. 5), Italy 6.9-fold (62 vs. 9), Russia 6.5-fold (13 vs. 2), Pakistan 5.7-fold (17 vs. 3), South Korea 5.2-fold (26 vs. 5). Detailed numerical data are in Supplementary Table 3.

Across countries with more than 50,000 authors with >=5 full articles when all scientific fields were considered, the highest proportions of authors with extreme publishing behavior among all authors were seen in Switzerland (823 of 81,045, 1.0%), Germany (1,791 of 370,960, 0.5%), and Italy (1,200 of 249,100, 0.5%). However, this was driven by the very strong participation of these countries in Physics multi-authored work. In the non-Physics group, among countries with more than 5,000 authors with >=5 full articles, the highest proportion of authors with extreme publishing behavior among all authors were seen in Saudi Arabia (98 of 27,588 authors, 0.36%) followed by Iraq (13 of 10,485, 0.12%), Malaysia (52 of 43,918, 0.12%), United Arab Emirates (9 of 8,059, 0.11%), Philippines (6 of 5,531, 0.11%), and Pakistan (33 of 32,529, 0.10%).

### Extreme publishing behavior across science: scientific field-level analysis

Extreme publishing behavior cluster more heavily in some scientific fields and subfields, while a few fields have still not witnessed this phenomenon. Figure 2 shows the share of each scientific field (excluding the field of Physics & Astronomy) in the total number of authors with extreme publishing behavior during 2000-2022. The patterns of clustering are similar for hyperprolific, almost hyperprolific and extreme publishing authors (detailed numerical data appear in Supplementary Table 4).

**Figure 2.**
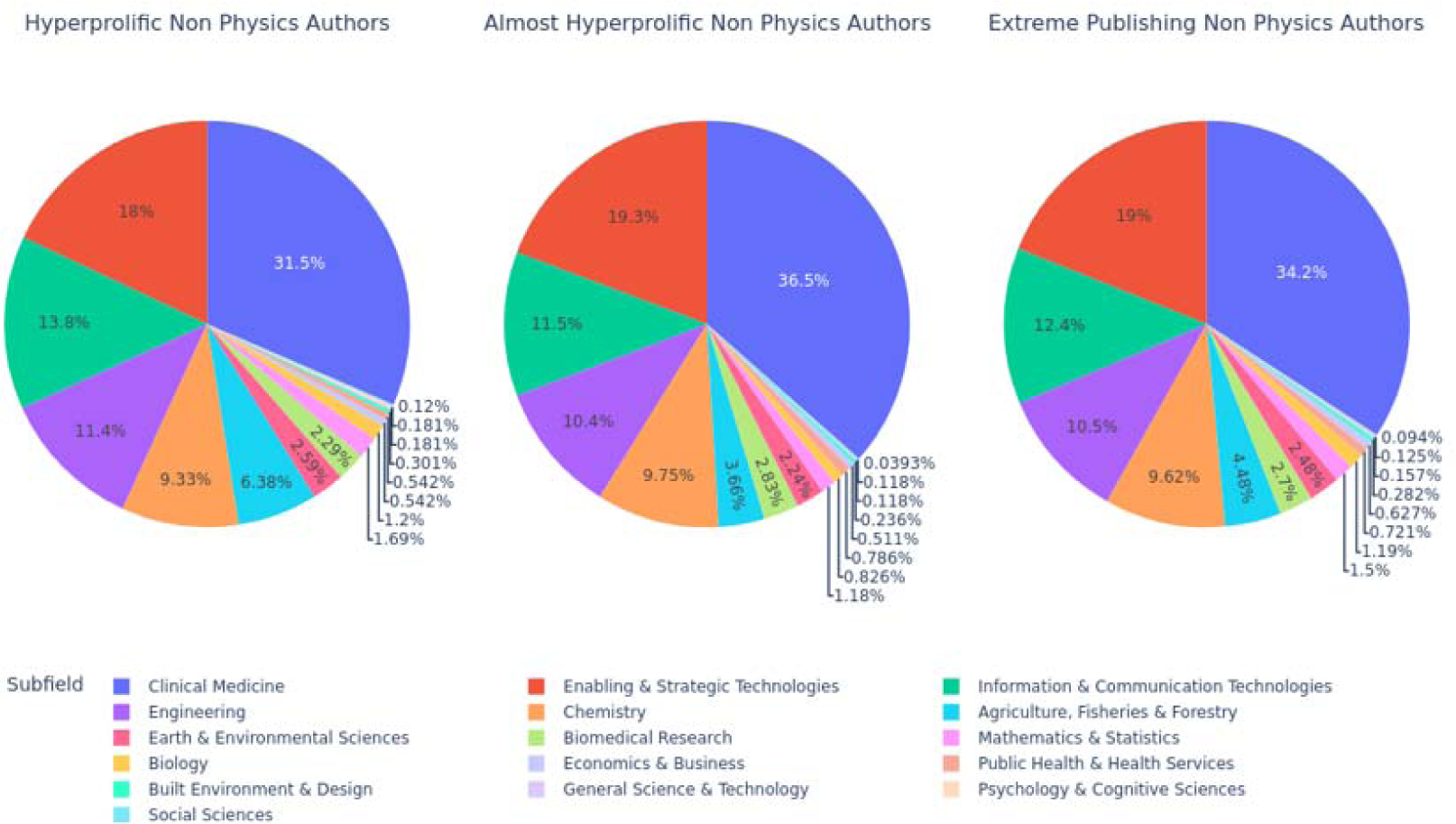
Pie charts of the number of hyperprolific, almost hyperprolific, and overall extreme publishing (sum of hyperprolific and almost hyperprolific) authors in non-Physics disciplines according to main scientific field for the entire period 2000-2022.

Among non-Physics fields, the scientific fields with the highest concentration of extreme publishing authors in 2022 were Clinical Medicine (n=678), Enabling & Strategic Technologies (n=327), Information & Communication Technologies (n=283), Engineering (n=168), Agriculture, Fisheries & Forestry (n=146), and Chemistry (n=140). There were modest numbers of extreme publishing authors that year in Earth & Environmental Sciences (n=57), Mathematics & Statistics (n=43), Biomedical Research (n=37), Biology (n=26), Economics & Business (n=17), and Public Health & Health Services (n=10), few extreme publishing authors in Built Environment & Design (n=6) and Psychology & Cognitive Sciences (n=2), and no extreme publishing authors in 5 fields (Social Sciences, Communication & Textual Studies, Historical Studies, Philosophy & Theology, Visual & Performing Arts). Compared with 2016, most fields saw a 2- to 4-fold increase in the number of extreme publishing authors in 2022, but there was a more dramatic increase in Agriculture, Fisheries & Forestry 14.6-fold (146 versus 10), Biology 13-fold (26 vs. 2), and Mathematics and Statistics 6.1-fold (43 vs. 7), while Economics & Business, Built Environment & Design and Psychology & Cognitive Sciences had no extreme publishing authors in 2016.

Given that the majority of extreme publishing authors in 2022 excluding Physics were in Clinical Medicine, we also examined whether specific subfields within this field had more major increases between 2016 and 2022. The highest fold-increases were seen in Complementary & Alternative Medicine (10.3-fold), Tropical Medicine (5.5-fold), Dentistry (4.5-fold), Immunology (4.2-fold), and Pharmacology & Pharmacy (3.8-fold). We also examined the subfields of Agriculture and Biology that had the most dramatic fold-increases between 2016 and 2022. The highest-fold increases were seen in Plant Biology & Botany (16 versus 1, 16-fold), Food science (78 versus 5, 15.6-fold), Fisheries (14 versus 1, 14-fold), Agronomy & Agriculture (13 versus 1, 13-fold) and Dairy & Animal Science (24 versus 2, 12-fold).

In terms of representation of extreme publishing authors among all authors with >=5 full articles in the field during 2000-2022, the highest proportions outside of Physics were seen in in Enabling and Strategic Technologies (605 of 685703 authors, 0.12%), Information and Communication Technologies (396 of 627,550, 0.08%), Chemistry (307 of 546,679, 0.08%), Engineering (334 of 456,772, 0.07%), and Agriculture, Fisheries and Forestry (143 of 211,946, 0.07%).

### Ranking for citation impact

Authors with extreme publishing behavior were a lion’s share of the authors in the highest ranks of citation impact, when this was quantified with raw total citations, but this dominance largely disappeared when a composite citation indicator was used to account also for co-authorship (Figure 3). Based on raw citation counts, authors with extreme publishing behavior accounted for 4,360 of the top-10,000 most-cited authors across science. In the Physics group, extreme publishing authors accounted for 3,336 /17,768 (18.8%) of the top-2% (with and/or without self-citations counted) according to raw citation counts but only 576/17,578 (3.28%) of the top-2% according to the composite citation indicator. In non-Physics fields, extreme publishing authors accounted for 2,402/184,391(1.30%) of the top-2% by raw citation counts and 2,139/184,113 (1.16%) of the top-2% according to the composite citation indicator. As shown in Figure 3, the large majority of extreme publishing authors across science excluding Physics have very prominent ranking even with the composite citation indicator (67.0% were in the top-2%), while most extreme publishing authors in Physics have very modest ranks with the composite citation indicator (4.56% are in the top-2%).

**Figure 3.**
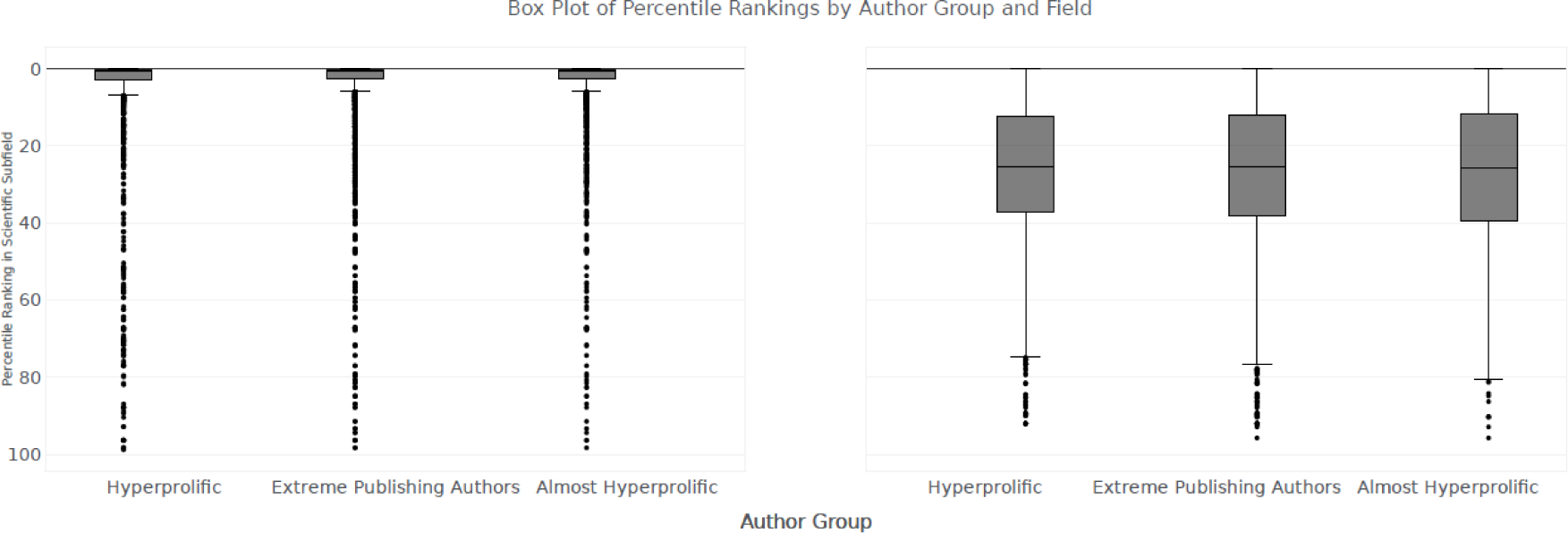
Boxplots of rankings based on composite citation indicator (all career-long, including self-citations) for hyperprolific, almost hyperprolific, and overall extreme publishing (sum of hyperprolific and almost hyperprolific) authors, separately for Physics and for all other scientific fields excluding Physics.

## CONCLUSIONS

The current analysis of the Scopus database has documented a massive increase in the number of authors who exhibit extreme publishing behavior in recent years in fields outside the discipline of nuclear and particle physics that has been well-known to operate with large-scale collaborations resulting in massive co-authorships of published articles. In fact, as the output of EONR overall and in terms of massively co-authored articles has recently declined, by 2022 the number of authors with extreme publishing behavior in fields outside of Physics has almost matched the number of such authors in Physics. Overall, during 2000-2022, more than 3,000 authors outside of Physics (and more than 12,000 in Physics) have had at least one calendar year where they published more than 1 full article every 6 days. Most of them had at least one calendar year where they published more than 1 full article every 5 days. Outside of Physics, the increase in the number of extreme publishing authors seems to have accelerated during 2016-2022 with a >3-fold increase in this period. However, some countries and some fields have witnessed a far more marked increase in extreme publishing behavior.

China has remained the country with the highest number of hyperprolific, almost hyperprolific, and overall extreme publishing authors for many consecutive years now. Overall, the ascendancy of China among authors with extreme publishing behavior may reflect the use of policies in China that placed emphasis in promoting publishing more, with major financial records (Xu et al., 2021; Quan et al., 2017). These policies have been heavily criticized and some of them have been reverted (Wang et al., 2021)]. Regardless, China is currently publishing more scientific articles than any other country (Xie & Freeman, 2019).

Eight countries had very large relative increases in the number of extreme publishing authors, amounting to 5-19-fold increases between 2016 and 2022. The advent of Saudi Arabia among the affiliation of extreme publishing authors may be due to the strong financial incentives offered by Saudi institutions. In Scopus, this reflects almost entirely local Saudi authors who develop extreme publishing behavior, not the listing of Saudi institutions by authors working mostly in other countries in the Clarivate Highly Cited Researchers database (Bhatacharjee, 2011; Catanzaro, 2023). Spain, Italy, and South Korea have also seen a spectacular increase in the number of hyperpprolific authors. It is unclear to which extent this may be related to specific national and/or university/institutional policies that favor raw increase in article numbers and English-language international publications over local language publications. Thailand, India, Russia and Pakistan have also witnessed sharp increases in extreme publishing author counts, even if the numbers are still modest in absolute terms when seen against their large populations. Excluding Physics, the highest presence of extreme publishing authors after adjusting for the total number of authors in each country is seen in Muslim countries (Saudi Arabia, Iraq, United Arab Emirates, Pakistan) and in Malaysia and Philippines. One may speculate whether geopolitical reasons, developing country economics, incentive systems, and local academic microenvironments may be promoting massive authorship practices in some scientists in these countries.

The majority of authors with extreme publishing behavior outside of Physics operate within the broad field of Clinical Medicine. This is not surprising, given that about one in three authors across science belong to this field. Nevertheless, some particular subfields seem to have a more major acceleration of the phenomenon. Agriculture, Fisheries & Forestry, Biology, and Mathematics and Statistics have also witnessed extreme relative increases in the number of extreme publishing authors in recent years. It is possible that these notorious increases reflect specific niches where extreme publishing behavior has become established and is adopted by several scientists working in these niches. Authorship practices may have become more lax in these niches, new norms of co-authorship may have evolved or unethical practices such as paper mills may have infiltrated these fields.

In our analysis, we made no effort to identify whether authors with extreme publishing behavior fulfill the typically required authorship criteria (e.g. Vancouver). However, based on previous survey results (Ioannidis et al., 2018), it is likely that many, if not most, of these authors do not routinely follow Vancouver criteria. Moreover, we made no effort to identify if some of these authors are associated with overtly unethical practices such as paper mills (Else & van Noorden, 2021; Christopher, 2021) or citation cartels (citation farms) (Fister et al. 2016). These characterizations would require in-depth evaluations of the CVs of single authors and meticulous investigative work. The Scopus data that we have made available may facilitate such efforts in the future.

Regardless of the exact of mix of genuinely high output, spurious authorship standards, and outright unethical research practices (Ioannidis & Maniadis, 2024), extreme publishing authors appear to enjoy high success in terms in citation impact, especially when raw citation counts are considered. 44% of the most-cited authors across science in terms of raw citations are extreme publishing authors. This suggests that counting citations without adjusting for co-authorship patterns may be highly problematic. Using a composite citation indicator that tries to correct for co-authorship and author position patterns, most of the Physics extreme publishing authors no longer rank very highly. Among non-Physics extreme publishing authors, the vast majority still reach the very top ranks of citation impact, even with these adjustments. This means that besides the sheer volume of published items, they often have influential position placements such as last author. This is a typical situation in some fields such as Clinical Medicine, where department leaders acquire the senior author spot, often with questionable research contributions or even overt gift authorship (Al-Herz et al., 2014; Kovac, 2013). While authors with extreme publishing behavior are a very small percentage of the scientific workforce, they have a substantial share among the ranks of the most-cited scientists. Given this high visibility and perceived impact, it is likely that many of them may also exert high influence in their environments and shape the course of science in their institutions and in their fields. This may make extreme publishing behavior not only legitimized but also highly coveted by other scientists in the same environment, propagating the further growth of the extreme publishing authors’ cohort.

Our work has some limitations. First, the identification of extreme publishing authors may be affected by errors in Scopus. Scopus author profile quality on precision (collecting the right publications for an author) and recall (collecting all publications for the author in the database in the profile) is continuously monitored using a gold set. In previous work done 7 years ago (Ioannidis et al., 2018), we had excluded authors with Chinese names, because at that time there was uncertainty about precision and recall for authors from Eastern Asia. Currently, the average overall precision reported by Elsevier by October 2023, is 96.6% with a recall of 92.1%. Specifically for authors in the gold set publishing in Eastern Asia, the precision is 97.1% and the recall 91.2% versus precision of 97.9% and a recall of 89.8% for authors publishing in North America.

Second, there is no absolute cut-off for the number of annually published articles that may be too much. Nevertheless, the use of two different thresholds gives qualitatively similar patterns for the features of extreme publishing behavior. Third, some journals are not covered by Scopus. Hence, the number of publications for some authors may be undercounted and the number of extreme publishing authors may be even larger than estimated here. Fourth, we did not account for other aspects of author output besides full articles. While journal-published items that are not full articles typically require far less effort to generate and publish, books may require major effort. It would be interesting to evaluate in the future also the presence of prolific book authors in databases that index published books.

Allowing for these caveats, our analyses show a major increase in authors with extreme publishing behavior across almost all scientific fields with multiple countries and fields and subfields leading this phenomenon. While some very talented, outstanding scientists may be included in the extreme publishing group, spurious and unethical behaviors may also abound. With the advent of mega-journals publishing many thousands of articles every year (Ioannidis et al., 2023), of artificial intelligence that may further facilitate writing articles (Flanagin, et al., 2023) and with peer-review having major limitations, the increase in extreme publishing authorship patterns is likely to continue in the future. Counting the number of published articles and using gamed citation metrics have acquired major influence in most scientific environments and they are often misused despite the availability of guidance for their more proper use (Hicks et al, 2015). Some authors have even argued that there should be a limit to the number of pages/articles a scientist can publish (Martinson, 2017). However, this may be a bad idea as it may further exaggerate publication bias and other selection biases. Instead, it may be more realistic and appropriate to monitor extreme publishing behaviors in centralized, standardized databases (Ioannidis & Maniadis, 2023), as we have done here. This monitoring should allow careful in-depth assessments of extreme patterns for single authors, teams, institutions, and countries.

## Supporting information

none

## Acknowledgments

A pre-print of this work has been deposited to bioRxiv, https://doi.org/10.1101/2023.11.23.568476

## Data sharing

All key data are in the manuscript and its supplementary files. More detailed data on the 3,191 extreme publishing authors in non-Physics scientific fields and on the 12,624 extreme publishing authors in Physics are available in https://elsevier.digitalcommonsdata.com/datasets/kmyvjk3xmd/2.

## Contributions

JPAI had the original idea and wrote the first draft of the article. TAC analyzed the data with contributions also from JPAI and JB. All authors discussed iterations of the protocol, interpreted the data and contributed writing the article and approved the final version. JPAI is guarantor.

## REFERENCES

Al-Herz W, Haider H, Al-Bahhar M & Sadeq A (2014). Honorary authorship in biomedical journals: how common is it and why does it exist? J Med Ethics. 40(5), 346–8.

Andersen LB, & Pallesen T (2008). “Not Just for the Money?” How Financial Incentives Affect the Number of Publications at Danish Research Institutions. International Public Management Journal 11, 28–47.

Archambault E, Beauchesne OH, & Caruso J (2011). “Towards a multilingual, comprehensive and open scientific journal ontology” in Proceedings of the 13th International Conference of the International Society for Scientometrics and Informetrics (ISSI), Durban, South Africa, B. Noyons, P. Ngulube, J. Leta, Eds. pp. 66–77.

Baas J, Schotten M, Plume M, Côté G, & Karimi R (2020). Scopus as a curated, high-quality bibliometric data source for academic research in quantitative science studies. Quant Sci Stud. 1, 377–386.

Bhattacharjee Y (2011). Saudi universities offer cash in exchange for academic prestige. Science 334(6061), 1344–1345.

Catanzaro M (2023). Saudi universities entice top scientists to switch affiliations-sometimes with cash. Nature. 617(7961), 446–447.

Chapman CA, Bicca-Marques JC, Calvignac-Spencer S, Fan P, Fashing PJ, Gogarten J, et al. (2019). Games academics play and their consequences: how authorship, h-index and journal impact factors are shaping the future of academia. Proc R Soc B. 286, 20192047.

Christopher J (2021). The raw truth about paper mills. FEBS letters. 595(13), 1751–1757.

Else H & Van Noorden R (2021). The fight against fake-paper factories that churn out sham science. Nature 591, 516–9.

Hicks D, Wouters P, Waltman L, De Rijcke S & Rafols I (2015). Bibliometrics: the Leiden Manifesto for research metrics. Nature 520 (7548), 429–431.

Fister I Jr, Fister I, & Perc M (2016). Towards the discovery of citation cartels in citation networks. Front Phys 4:00049.

Flanagin A, Bibbins-Domingo K, Berkwits M, & Christiansen SL (2023). Nonhuman “Authors” and Implications for the Integrity of Scientific Publication and Medical Knowledge. JAMA. 329, 637–639.

Fontanarosa P, Bauchner H, & Flanagin A (2017). Authorship and Team Science. JAMA. 318(24), 2433–2437.

Hosseini, M., Lewis, J., Zwart, H. et al. (2022) An ethical exploration of increased average number of authors per publication. Sci Eng Ethics 28, 25.

Ioannidis JPA, Klavans R & Boyack KW (2018). Thousands of scientists publish a paper every five days. Nature. 561(7722), 167–169.

Ioannidis JPA, Baas J, Klavans R, & Boyack KW (2019). A standardized citation metrics author database annotated for scientific field. PLoS Biol. 17(8), e3000384.

Ioannidis JP, Klavans R & Boyack KW (2016). Multiple Citation Indicators and Their Composite across Scientific Disciplines. PLoS Biol. 14(7):e1002501.

Ioannidis JPA & Maniadis Z (2024). Quantitative research assessment: metrics against gamed metrics. Intern Emerg Med. doi: 10.1007/s11739-023-03447-w.

Ioannidis JPA, Pezzullo AM & Boccia S (2023). The Rapid Growth of Mega-Journals: Threats and Opportunities. JAMA. 329(15), 1253–1254.

Ioannidis JP & Maniadis Z (2023). In defense of quantitative researcher assessments. PLoS Biol. (in press)

Kim DH & Bak H-J (2016). How Do Scientists Respond to Performance-Based Incentives? Evidence From South Korea. International Public Management Journal 19, 31–52.

Kovacs J (2013). Honorary authorship epidemic in scholarly publications? How the current use of citation-based evaluative metrics make (pseudo)honorary authors from honest contributors of every multi-author article. J Med Ethics. 39(8), 509–12.

Martinson B (2017). Give researchers a lifetime word limit. Nature 550, 303.

Papatheodorou SI, Trikalinos TA & Ioannidis JP (2008). Inflated numbers of authors over time have not been just due to increasing research complexity. J Clin Epidemiol. 61(6), 546–51.

Quan W, Chen B, & Shu F (2017). Publish or impoverish: An investigation of the monetary reward system of science in China (1999-2016). Aslib Journal of Information Management, 69, 486–502.

Xie Q & Freeman RB (2019). Bigger than you thought: China’s contribution to scientific publications and its impact on the global economy. China & World Economy 27(1), 1–27.

Xu X, Rose H & Oancea A (2021). Incentivising international publications: institutional policymaking in Chinese higher education, Studies in Higher Education 46, 1132–1145.

Wager E, Singhvi S, & Kleinert S (2015). Too much of a good thing? An observational study of prolific authors. PeerJ. 3, e1154.

Wang J, Halffman W & Zwart H (2021). The Chinese scientific publication system: Specific features, specific challenges. Learned Publishing 34, 105–115.

