## Supplementary material for "Evolving patterns of extreme publishing behavior across science": none

Supplementary Table 1 and Supplementary Figure 1. Number of full papers overall and with >100, >500, and >1000 authors published with an affiliation from the European Organization for Nuclear Research

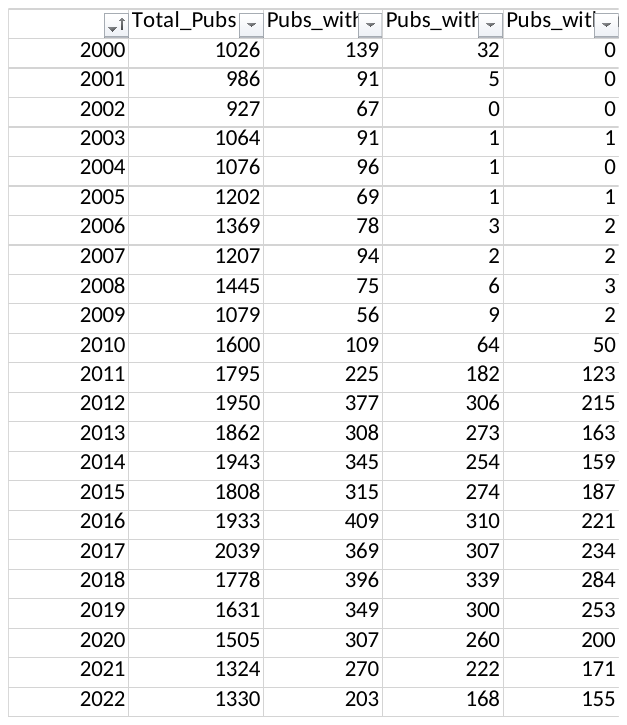

Pubs with Author Count for EONR

Publication Count

| Year | Total_Pubs | Pubs_with_more_than_100_authors | Pubs_with_more_than_500_authors | Pubs_with_more_than_1000_authors |
| --- | --- | --- | --- | --- |
| 2000 | 1026 | 139 | 32 | 0 |
| 2001 | 986 | 91 | 5 | 0 |
| 2002 | 927 | 67 | 0 | 0 |
| 2003 | 1064 | 91 | 1 | 1 |
| 2004 | 1076 | 96 | 1 | 0 |
| 2005 | 1202 | 69 | 1 | 1 |
| 2006 | 1369 | 78 | 3 | 2 |
| 2007 | 1207 | 94 | 2 | 2 |
| 2008 | 1445 | 75 | 6 | 3 |
| 2009 | 1079 | 56 | 9 | 2 |
| 2010 | 1600 | 109 | 64 | 50 |
| 2011 | 1795 | 225 | 182 | 123 |
| 2012 | 1950 | 377 | 306 | 215 |
| 2013 | 1862 | 308 | 273 | 163 |
| 2014 | 1943 | 345 | 254 | 159 |
| 2015 | 1808 | 315 | 274 | 187 |
| 2016 | 1933 | 409 | 310 | 221 |
| 2017 | 2039 | 369 | 307 | 234 |
| 2018 | 1778 | 396 | 339 | 284 |
| 2019 | 1631 | 349 | 300 | 253 |
| 2020 | 1505 | 307 | 260 | 200 |
| 2021 | 1324 | 270 | 222 | 171 |
| 2022 | 1330 | 203 | 168 | 155 |

Supplementary Table 2. Number of HP, AHP, and overall EP (sum of HP and AHP) authors in each country during 2000-2022 for Physics and for non-Physics scientific fields

| cntry | Number of authors | Group | Rank |
| --- | --- | --- | --- |
| chn | 433 | HP_Non_Physics_Authors | 1 |
| usa | 185 | HP_Non_Physics_Authors | 2 |
| deu | 82 | HP_Non_Physics_Authors | 3 |
| others | 80 | HP_Non_Physics_Authors | 4 |
| gbr | 77 | HP_Non_Physics_Authors | 5 |
| ind | 65 | HP_Non_Physics_Authors | 6 |
| ita | 62 | HP_Non_Physics_Authors | 7 |
| jpn | 55 | HP_Non_Physics_Authors | 8 |
| sau | 54 | HP_Non_Physics_Authors | 9 |
| aus | 52 | HP_Non_Physics_Authors | 10 |
| kor | 31 | HP_Non_Physics_Authors | 11 |
| twn | 29 | HP_Non_Physics_Authors | 12 |
| mys | 27 | HP_Non_Physics_Authors | 13 |
| irn | 27 | HP_Non_Physics_Authors | 13 |
| fra | 26 | HP_Non_Physics_Authors | 15 |
| nld | 25 | HP_Non_Physics_Authors | 16 |
| can | 25 | HP_Non_Physics_Authors | 16 |
| esp | 24 | HP_Non_Physics_Authors | 18 |
| pak | 19 | HP_Non_Physics_Authors | 19 |
| dnk | 18 | HP_Non_Physics_Authors | 20 |
| prt | 14 | HP_Non_Physics_Authors | 21 |
| tha | 14 | HP_Non_Physics_Authors | 21 |
| sgp | 13 | HP_Non_Physics_Authors | 23 |
| egy | 13 | HP_Non_Physics_Authors | 23 |
| zaf | 13 | HP_Non_Physics_Authors | 23 |
| che | 11 | HP_Non_Physics_Authors | 26 |
| grc | 11 | HP_Non_Physics_Authors | 26 |
| rus | 10 | HP_Non_Physics_Authors | 28 |
| bra | 9 | HP_Non_Physics_Authors | 29 |
| tur | 9 | HP_Non_Physics_Authors | 29 |
| irq | 8 | HP_Non_Physics_Authors | 31 |
| hkg | 8 | HP_Non_Physics_Authors | 31 |
| isr | 8 | HP_Non_Physics_Authors | 31 |
| nor | 7 | HP_Non_Physics_Authors | 34 |
| idn | 7 | HP_Non_Physics_Authors | 34 |
| bel | 7 | HP_Non_Physics_Authors | 34 |
| pol | 6 | HP_Non_Physics_Authors | 37 |
| bgd | 6 | HP_Non_Physics_Authors | 37 |
| are | 6 | HP_Non_Physics_Authors | 37 |
| rou | 5 | HP_Non_Physics_Authors | 40 |
| fin | 5 | HP_Non_Physics_Authors | 40 |
| swe | 5 | HP_Non_Physics_Authors | 40 |
| cze | 5 | HP_Non_Physics_Authors | 40 |
| vnm | 4 | HP_Non_Physics_Authors | 44 |
| aut | 4 | HP_Non_Physics_Authors | 44 |
| chl | 3 | HP_Non_Physics_Authors | 46 |
| irl | 3 | HP_Non_Physics_Authors | 46 |
| bgr | 3 | HP_Non_Physics_Authors | 46 |
| jor | 3 | HP_Non_Physics_Authors | 46 |
| phl | 3 | HP_Non_Physics_Authors | 46 |
| lux | 2 | HP_Non_Physics_Authors | 51 |
| srb | 2 | HP_Non_Physics_Authors | 51 |
| qat | 2 | HP_Non_Physics_Authors | 51 |
| svk | 2 | HP_Non_Physics_Authors | 51 |
| cyp | 2 | HP_Non_Physics_Authors | 51 |
| nga | 2 | HP_Non_Physics_Authors | 51 |
| mar | 2 | HP_Non_Physics_Authors | 51 |
| nzl | 2 | HP_Non_Physics_Authors | 51 |
| lbn | 2 | HP_Non_Physics_Authors | 51 |
| mac | 2 | HP_Non_Physics_Authors | 51 |
| mex | 2 | HP_Non_Physics_Authors | 51 |
| kwt | 1 | HP_Non_Physics_Authors | 62 |
| geo | 1 | HP_Non_Physics_Authors | 62 |
| per | 1 | HP_Non_Physics_Authors | 62 |
| omn | 1 | HP_Non_Physics_Authors | 62 |
| dji | 1 | HP_Non_Physics_Authors | 62 |
| ukr | 1 | HP_Non_Physics_Authors | 62 |
| brn | 1 | HP_Non_Physics_Authors | 62 |
| isl | 1 | HP_Non_Physics_Authors | 62 |
| grd | 1 | HP_Non_Physics_Authors | 62 |
| gha | 1 | HP_Non_Physics_Authors | 62 |
| hun | 1 | HP_Non_Physics_Authors | 62 |
| mus | 1 | HP_Non_Physics_Authors | 62 |
| syr | 1 | HP_Non_Physics_Authors | 62 |
| pse | 1 | HP_Non_Physics_Authors | 62 |
| eth | 1 | HP_Non_Physics_Authors | 62 |
| ecu | 1 | HP_Non_Physics_Authors | 62 |
| uzb | 1 | HP_Non_Physics_Authors | 62 |
| ltu | 1 | HP_Non_Physics_Authors | 62 |
| tun | 1 | HP_Non_Physics_Authors | 62 |
| kaz | 1 | HP_Non_Physics_Authors | 62 |
| usa | 2587 | HP_Physics_Authors | 1 |
| deu | 1346 | HP_Physics_Authors | 2 |
| ita | 892 | HP_Physics_Authors | 3 |
| gbr | 815 | HP_Physics_Authors | 4 |
| che | 697 | HP_Physics_Authors | 5 |
| fra | 482 | HP_Physics_Authors | 6 |
| rus | 399 | HP_Physics_Authors | 7 |
| chn | 378 | HP_Physics_Authors | 8 |
| jpn | 257 | HP_Physics_Authors | 9 |
| esp | 224 | HP_Physics_Authors | 10 |
| can | 218 | HP_Physics_Authors | 11 |
| bel | 155 | HP_Physics_Authors | 12 |
| ind | 123 | HP_Physics_Authors | 13 |
| cze | 119 | HP_Physics_Authors | 14 |
| nld | 118 | HP_Physics_Authors | 15 |
| swe | 115 | HP_Physics_Authors | 16 |
| pol | 110 | HP_Physics_Authors | 17 |
| kor | 109 | HP_Physics_Authors | 18 |
| tur | 97 | HP_Physics_Authors | 19 |
| bra | 88 | HP_Physics_Authors | 20 |
| grc | 87 | HP_Physics_Authors | 21 |
| isr | 71 | HP_Physics_Authors | 22 |
| aus | 68 | HP_Physics_Authors | 23 |
| prt | 65 | HP_Physics_Authors | 24 |
| twn | 64 | HP_Physics_Authors | 25 |
| aut | 57 | HP_Physics_Authors | 26 |
| hun | 48 | HP_Physics_Authors | 27 |
| nor | 46 | HP_Physics_Authors | 28 |
| rou | 44 | HP_Physics_Authors | 29 |
| dnk | 38 | HP_Physics_Authors | 30 |
| fin | 34 | HP_Physics_Authors | 31 |
| mex | 33 | HP_Physics_Authors | 32 |
| zaf | 27 | HP_Physics_Authors | 33 |
| bgr | 25 | HP_Physics_Authors | 34 |
| srb | 23 | HP_Physics_Authors | 35 |
| hrv | 23 | HP_Physics_Authors | 35 |
| col | 22 | HP_Physics_Authors | 37 |
| mar | 22 | HP_Physics_Authors | 37 |
| pak | 22 | HP_Physics_Authors | 37 |
| sau | 20 | HP_Physics_Authors | 40 |
| hkg | 20 | HP_Physics_Authors | 40 |
| svk | 20 | HP_Physics_Authors | 40 |
| irn | 19 | HP_Physics_Authors | 43 |
| chl | 18 | HP_Physics_Authors | 44 |
| arg | 17 | HP_Physics_Authors | 45 |
| mys | 17 | HP_Physics_Authors | 45 |
| cyp | 16 | HP_Physics_Authors | 47 |
| svn | 16 | HP_Physics_Authors | 47 |
| geo | 14 | HP_Physics_Authors | 49 |
| blr | 14 | HP_Physics_Authors | 49 |
| est | 13 | HP_Physics_Authors | 51 |
| tha | 9 | HP_Physics_Authors | 52 |
| ltu | 9 | HP_Physics_Authors | 52 |
| ukr | 8 | HP_Physics_Authors | 54 |
| nzl | 8 | HP_Physics_Authors | 54 |
| arm | 6 | HP_Physics_Authors | 56 |
| egy | 6 | HP_Physics_Authors | 56 |
| irl | 5 | HP_Physics_Authors | 58 |
| pri | 4 | HP_Physics_Authors | 59 |
| idn | 4 | HP_Physics_Authors | 59 |
| aze | 4 | HP_Physics_Authors | 59 |
| sgp | 3 | HP_Physics_Authors | 62 |
| vnm | 3 | HP_Physics_Authors | 62 |
| qat | 2 | HP_Physics_Authors | 64 |
| dza | 2 | HP_Physics_Authors | 64 |
| others | 2 | HP_Physics_Authors | 64 |
| ecu | 2 | HP_Physics_Authors | 64 |
| lka | 2 | HP_Physics_Authors | 64 |
| irq | 1 | HP_Physics_Authors | 69 |
| per | 1 | HP_Physics_Authors | 69 |
| lva | 1 | HP_Physics_Authors | 69 |
| omn | 1 | HP_Physics_Authors | 69 |
| mne | 1 | HP_Physics_Authors | 69 |
| bgd | 1 | HP_Physics_Authors | 69 |
| pse | 1 | HP_Physics_Authors | 69 |
| are | 1 | HP_Physics_Authors | 69 |
| jor | 1 | HP_Physics_Authors | 69 |
| kaz | 1 | HP_Physics_Authors | 69 |
| chn | 711 | AHP_Non_Physics_Authors | 1 |
| usa | 339 | AHP_Non_Physics_Authors | 2 |
| deu | 166 | AHP_Non_Physics_Authors | 3 |
| jpn | 120 | AHP_Non_Physics_Authors | 4 |
| gbr | 114 | AHP_Non_Physics_Authors | 5 |
| ita | 99 | AHP_Non_Physics_Authors | 6 |
| aus | 83 | AHP_Non_Physics_Authors | 7 |
| ind | 63 | AHP_Non_Physics_Authors | 8 |
| sau | 60 | AHP_Non_Physics_Authors | 9 |
| can | 55 | AHP_Non_Physics_Authors | 10 |
| irn | 50 | AHP_Non_Physics_Authors | 11 |
| others | 45 | AHP_Non_Physics_Authors | 12 |
| nld | 43 | AHP_Non_Physics_Authors | 13 |
| esp | 40 | AHP_Non_Physics_Authors | 14 |
| mys | 39 | AHP_Non_Physics_Authors | 15 |
| kor | 38 | AHP_Non_Physics_Authors | 16 |
| fra | 29 | AHP_Non_Physics_Authors | 17 |
| twn | 29 | AHP_Non_Physics_Authors | 17 |
| dnk | 25 | AHP_Non_Physics_Authors | 19 |
| rus | 24 | AHP_Non_Physics_Authors | 20 |
| sgp | 22 | AHP_Non_Physics_Authors | 21 |
| pak | 22 | AHP_Non_Physics_Authors | 21 |
| che | 21 | AHP_Non_Physics_Authors | 23 |
| hkg | 20 | AHP_Non_Physics_Authors | 24 |
| grc | 19 | AHP_Non_Physics_Authors | 25 |
| bra | 18 | AHP_Non_Physics_Authors | 26 |
| tur | 17 | AHP_Non_Physics_Authors | 27 |
| idn | 16 | AHP_Non_Physics_Authors | 28 |
| tha | 16 | AHP_Non_Physics_Authors | 28 |
| bel | 16 | AHP_Non_Physics_Authors | 28 |
| prt | 13 | AHP_Non_Physics_Authors | 31 |
| fin | 11 | AHP_Non_Physics_Authors | 32 |
| swe | 11 | AHP_Non_Physics_Authors | 32 |
| cze | 11 | AHP_Non_Physics_Authors | 32 |
| zaf | 11 | AHP_Non_Physics_Authors | 32 |
| egy | 10 | AHP_Non_Physics_Authors | 36 |
| vnm | 9 | AHP_Non_Physics_Authors | 37 |
| pol | 8 | AHP_Non_Physics_Authors | 38 |
| isr | 8 | AHP_Non_Physics_Authors | 38 |
| rou | 7 | AHP_Non_Physics_Authors | 40 |
| irq | 6 | AHP_Non_Physics_Authors | 41 |
| mac | 6 | AHP_Non_Physics_Authors | 41 |
| col | 5 | AHP_Non_Physics_Authors | 43 |
| are | 5 | AHP_Non_Physics_Authors | 43 |
| qat | 4 | AHP_Non_Physics_Authors | 45 |
| bgd | 4 | AHP_Non_Physics_Authors | 45 |
| irl | 4 | AHP_Non_Physics_Authors | 45 |
| aut | 4 | AHP_Non_Physics_Authors | 45 |
| nor | 3 | AHP_Non_Physics_Authors | 49 |
| isl | 3 | AHP_Non_Physics_Authors | 49 |
| nzl | 3 | AHP_Non_Physics_Authors | 49 |
| lbn | 3 | AHP_Non_Physics_Authors | 49 |
| phl | 3 | AHP_Non_Physics_Authors | 49 |
| lux | 2 | AHP_Non_Physics_Authors | 54 |
| chl | 2 | AHP_Non_Physics_Authors | 54 |
| srb | 2 | AHP_Non_Physics_Authors | 54 |
| omn | 2 | AHP_Non_Physics_Authors | 54 |
| nga | 2 | AHP_Non_Physics_Authors | 54 |
| mar | 2 | AHP_Non_Physics_Authors | 54 |
| jor | 2 | AHP_Non_Physics_Authors | 54 |
| ltu | 2 | AHP_Non_Physics_Authors | 54 |
| tun | 2 | AHP_Non_Physics_Authors | 54 |
| kwt | 1 | AHP_Non_Physics_Authors | 63 |
| per | 1 | AHP_Non_Physics_Authors | 63 |
| lva | 1 | AHP_Non_Physics_Authors | 63 |
| dji | 1 | AHP_Non_Physics_Authors | 63 |
| brn | 1 | AHP_Non_Physics_Authors | 63 |
| cyp | 1 | AHP_Non_Physics_Authors | 63 |
| grd | 1 | AHP_Non_Physics_Authors | 63 |
| gha | 1 | AHP_Non_Physics_Authors | 63 |
| hun | 1 | AHP_Non_Physics_Authors | 63 |
| mus | 1 | AHP_Non_Physics_Authors | 63 |
| syr | 1 | AHP_Non_Physics_Authors | 63 |
| ken | 1 | AHP_Non_Physics_Authors | 63 |
| bgr | 1 | AHP_Non_Physics_Authors | 63 |
| mex | 1 | AHP_Non_Physics_Authors | 63 |
| usa | 1930 | AHP_Physics_Authors | 1 |
| deu | 964 | AHP_Physics_Authors | 2 |
| ita | 814 | AHP_Physics_Authors | 3 |
| gbr | 704 | AHP_Physics_Authors | 4 |
| chn | 590 | AHP_Physics_Authors | 5 |
| che | 541 | AHP_Physics_Authors | 6 |
| fra | 388 | AHP_Physics_Authors | 7 |
| rus | 338 | AHP_Physics_Authors | 8 |
| esp | 228 | AHP_Physics_Authors | 9 |
| jpn | 218 | AHP_Physics_Authors | 10 |
| can | 169 | AHP_Physics_Authors | 11 |
| ind | 103 | AHP_Physics_Authors | 12 |
| pol | 102 | AHP_Physics_Authors | 13 |
| bel | 94 | AHP_Physics_Authors | 14 |
| tur | 93 | AHP_Physics_Authors | 15 |
| kor | 93 | AHP_Physics_Authors | 15 |
| nld | 92 | AHP_Physics_Authors | 17 |
| bra | 87 | AHP_Physics_Authors | 18 |
| cze | 77 | AHP_Physics_Authors | 19 |
| swe | 76 | AHP_Physics_Authors | 20 |
| grc | 67 | AHP_Physics_Authors | 21 |
| isr | 56 | AHP_Physics_Authors | 22 |
| twn | 54 | AHP_Physics_Authors | 23 |
| aus | 46 | AHP_Physics_Authors | 24 |
| hun | 44 | AHP_Physics_Authors | 25 |
| prt | 39 | AHP_Physics_Authors | 26 |
| rou | 36 | AHP_Physics_Authors | 27 |
| aut | 36 | AHP_Physics_Authors | 27 |
| nor | 31 | AHP_Physics_Authors | 29 |
| fin | 30 | AHP_Physics_Authors | 30 |
| mex | 28 | AHP_Physics_Authors | 31 |
| pak | 25 | AHP_Physics_Authors | 32 |
| hrv | 24 | AHP_Physics_Authors | 33 |
| col | 23 | AHP_Physics_Authors | 34 |
| zaf | 23 | AHP_Physics_Authors | 34 |
| sau | 20 | AHP_Physics_Authors | 36 |
| mar | 20 | AHP_Physics_Authors | 36 |
| bgr | 19 | AHP_Physics_Authors | 38 |
| hkg | 18 | AHP_Physics_Authors | 39 |
| svn | 18 | AHP_Physics_Authors | 39 |
| dnk | 16 | AHP_Physics_Authors | 41 |
| srb | 15 | AHP_Physics_Authors | 42 |
| svk | 15 | AHP_Physics_Authors | 42 |
| arg | 14 | AHP_Physics_Authors | 44 |
| blr | 14 | AHP_Physics_Authors | 44 |
| irn | 14 | AHP_Physics_Authors | 44 |
| cyp | 13 | AHP_Physics_Authors | 47 |
| mys | 13 | AHP_Physics_Authors | 47 |
| geo | 12 | AHP_Physics_Authors | 49 |
| ltu | 9 | AHP_Physics_Authors | 50 |
| est | 9 | AHP_Physics_Authors | 50 |
| egy | 8 | AHP_Physics_Authors | 52 |
| lka | 8 | AHP_Physics_Authors | 52 |
| chl | 7 | AHP_Physics_Authors | 54 |
| ukr | 7 | AHP_Physics_Authors | 54 |
| tha | 7 | AHP_Physics_Authors | 54 |
| idn | 6 | AHP_Physics_Authors | 57 |
| sgp | 5 | AHP_Physics_Authors | 58 |
| others | 5 | AHP_Physics_Authors | 58 |
| irl | 4 | AHP_Physics_Authors | 60 |
| dza | 3 | AHP_Physics_Authors | 61 |
| vnm | 3 | AHP_Physics_Authors | 61 |
| arm | 3 | AHP_Physics_Authors | 61 |
| nzl | 3 | AHP_Physics_Authors | 61 |
| mng | 2 | AHP_Physics_Authors | 65 |
| lva | 2 | AHP_Physics_Authors | 65 |
| mne | 2 | AHP_Physics_Authors | 65 |
| ecu | 2 | AHP_Physics_Authors | 65 |
| kaz | 2 | AHP_Physics_Authors | 65 |
| pri | 1 | AHP_Physics_Authors | 70 |
| irq | 1 | AHP_Physics_Authors | 70 |
| omn | 1 | AHP_Physics_Authors | 70 |
| qat | 1 | AHP_Physics_Authors | 70 |
| aze | 1 | AHP_Physics_Authors | 70 |
| pse | 1 | AHP_Physics_Authors | 70 |
| are | 1 | AHP_Physics_Authors | 70 |
| usa | 2984 | HP_and_AHP_Physics_Authors | 1 |
| deu | 1603 | HP_and_AHP_Physics_Authors | 2 |
| ita | 1077 | HP_and_AHP_Physics_Authors | 3 |
| gbr | 985 | HP_and_AHP_Physics_Authors | 4 |
| che | 798 | HP_and_AHP_Physics_Authors | 5 |
| chn | 728 | HP_and_AHP_Physics_Authors | 6 |
| fra | 544 | HP_and_AHP_Physics_Authors | 7 |
| rus | 468 | HP_and_AHP_Physics_Authors | 8 |
| jpn | 349 | HP_and_AHP_Physics_Authors | 9 |
| esp | 277 | HP_and_AHP_Physics_Authors | 10 |
| can | 257 | HP_and_AHP_Physics_Authors | 11 |
| bel | 172 | HP_and_AHP_Physics_Authors | 12 |
| ind | 155 | HP_and_AHP_Physics_Authors | 13 |
| nld | 142 | HP_and_AHP_Physics_Authors | 14 |
| pol | 136 | HP_and_AHP_Physics_Authors | 15 |
| kor | 136 | HP_and_AHP_Physics_Authors | 15 |
| cze | 134 | HP_and_AHP_Physics_Authors | 17 |
| swe | 128 | HP_and_AHP_Physics_Authors | 18 |
| tur | 114 | HP_and_AHP_Physics_Authors | 19 |
| bra | 110 | HP_and_AHP_Physics_Authors | 20 |
| grc | 98 | HP_and_AHP_Physics_Authors | 21 |
| isr | 79 | HP_and_AHP_Physics_Authors | 22 |
| aus | 79 | HP_and_AHP_Physics_Authors | 22 |
| twn | 74 | HP_and_AHP_Physics_Authors | 24 |
| prt | 72 | HP_and_AHP_Physics_Authors | 25 |
| aut | 64 | HP_and_AHP_Physics_Authors | 26 |
| hun | 54 | HP_and_AHP_Physics_Authors | 27 |
| nor | 52 | HP_and_AHP_Physics_Authors | 28 |
| rou | 51 | HP_and_AHP_Physics_Authors | 29 |
| mex | 48 | HP_and_AHP_Physics_Authors | 30 |
| dnk | 39 | HP_and_AHP_Physics_Authors | 31 |
| fin | 36 | HP_and_AHP_Physics_Authors | 32 |
| pak | 34 | HP_and_AHP_Physics_Authors | 33 |
| sau | 33 | HP_and_AHP_Physics_Authors | 34 |
| hrv | 32 | HP_and_AHP_Physics_Authors | 35 |
| zaf | 32 | HP_and_AHP_Physics_Authors | 35 |
| bgr | 31 | HP_and_AHP_Physics_Authors | 37 |
| col | 30 | HP_and_AHP_Physics_Authors | 38 |
| mar | 28 | HP_and_AHP_Physics_Authors | 39 |
| hkg | 25 | HP_and_AHP_Physics_Authors | 40 |
| srb | 24 | HP_and_AHP_Physics_Authors | 41 |
| svk | 24 | HP_and_AHP_Physics_Authors | 41 |
| svn | 24 | HP_and_AHP_Physics_Authors | 41 |
| arg | 21 | HP_and_AHP_Physics_Authors | 44 |
| mys | 21 | HP_and_AHP_Physics_Authors | 44 |
| blr | 20 | HP_and_AHP_Physics_Authors | 46 |
| chl | 19 | HP_and_AHP_Physics_Authors | 47 |
| irn | 19 | HP_and_AHP_Physics_Authors | 47 |
| cyp | 16 | HP_and_AHP_Physics_Authors | 49 |
| geo | 14 | HP_and_AHP_Physics_Authors | 50 |
| est | 13 | HP_and_AHP_Physics_Authors | 51 |
| ltu | 11 | HP_and_AHP_Physics_Authors | 52 |
| egy | 10 | HP_and_AHP_Physics_Authors | 53 |
| tha | 10 | HP_and_AHP_Physics_Authors | 53 |
| ukr | 9 | HP_and_AHP_Physics_Authors | 55 |
| nzl | 9 | HP_and_AHP_Physics_Authors | 55 |
| idn | 8 | HP_and_AHP_Physics_Authors | 57 |
| lka | 8 | HP_and_AHP_Physics_Authors | 57 |
| sgp | 7 | HP_and_AHP_Physics_Authors | 59 |
| others | 6 | HP_and_AHP_Physics_Authors | 60 |
| arm | 6 | HP_and_AHP_Physics_Authors | 60 |
| irl | 6 | HP_and_AHP_Physics_Authors | 60 |
| pri | 4 | HP_and_AHP_Physics_Authors | 63 |
| dza | 4 | HP_and_AHP_Physics_Authors | 63 |
| vnm | 4 | HP_and_AHP_Physics_Authors | 63 |
| aze | 4 | HP_and_AHP_Physics_Authors | 63 |
| mng | 2 | HP_and_AHP_Physics_Authors | 67 |
| irq | 2 | HP_and_AHP_Physics_Authors | 67 |
| lva | 2 | HP_and_AHP_Physics_Authors | 67 |
| mne | 2 | HP_and_AHP_Physics_Authors | 67 |
| qat | 2 | HP_and_AHP_Physics_Authors | 67 |
| are | 2 | HP_and_AHP_Physics_Authors | 67 |
| ecu | 2 | HP_and_AHP_Physics_Authors | 67 |
| kaz | 2 | HP_and_AHP_Physics_Authors | 67 |
| per | 1 | HP_and_AHP_Physics_Authors | 75 |
| omn | 1 | HP_and_AHP_Physics_Authors | 75 |
| bgd | 1 | HP_and_AHP_Physics_Authors | 75 |
| pse | 1 | HP_and_AHP_Physics_Authors | 75 |
| jor | 1 | HP_and_AHP_Physics_Authors | 75 |
| chn | 846 | HP_and_AHP_Non_Physics_Authors | 1 |
| usa | 398 | HP_and_AHP_Non_Physics_Authors | 2 |
| deu | 188 | HP_and_AHP_Non_Physics_Authors | 3 |
| gbr | 135 | HP_and_AHP_Non_Physics_Authors | 4 |
| jpn | 134 | HP_and_AHP_Non_Physics_Authors | 5 |
| ita | 123 | HP_and_AHP_Non_Physics_Authors | 6 |
| others | 109 | HP_and_AHP_Non_Physics_Authors | 7 |
| aus | 104 | HP_and_AHP_Non_Physics_Authors | 8 |
| ind | 103 | HP_and_AHP_Non_Physics_Authors | 9 |
| sau | 98 | HP_and_AHP_Non_Physics_Authors | 10 |
| can | 59 | HP_and_AHP_Non_Physics_Authors | 11 |
| irn | 58 | HP_and_AHP_Non_Physics_Authors | 12 |
| kor | 56 | HP_and_AHP_Non_Physics_Authors | 13 |
| mys | 52 | HP_and_AHP_Non_Physics_Authors | 14 |
| nld | 48 | HP_and_AHP_Non_Physics_Authors | 15 |
| esp | 46 | HP_and_AHP_Non_Physics_Authors | 16 |
| fra | 41 | HP_and_AHP_Non_Physics_Authors | 17 |
| twn | 39 | HP_and_AHP_Non_Physics_Authors | 18 |
| pak | 33 | HP_and_AHP_Non_Physics_Authors | 19 |
| rus | 32 | HP_and_AHP_Non_Physics_Authors | 20 |
| dnk | 30 | HP_and_AHP_Non_Physics_Authors | 21 |
| sgp | 29 | HP_and_AHP_Non_Physics_Authors | 22 |
| che | 25 | HP_and_AHP_Non_Physics_Authors | 23 |
| tha | 25 | HP_and_AHP_Non_Physics_Authors | 23 |
| hkg | 24 | HP_and_AHP_Non_Physics_Authors | 25 |
| grc | 22 | HP_and_AHP_Non_Physics_Authors | 26 |
| idn | 21 | HP_and_AHP_Non_Physics_Authors | 27 |
| bra | 20 | HP_and_AHP_Non_Physics_Authors | 28 |
| tur | 20 | HP_and_AHP_Non_Physics_Authors | 28 |
| zaf | 20 | HP_and_AHP_Non_Physics_Authors | 28 |
| egy | 19 | HP_and_AHP_Non_Physics_Authors | 31 |
| bel | 19 | HP_and_AHP_Non_Physics_Authors | 31 |
| prt | 18 | HP_and_AHP_Non_Physics_Authors | 33 |
| swe | 14 | HP_and_AHP_Non_Physics_Authors | 34 |
| irq | 13 | HP_and_AHP_Non_Physics_Authors | 35 |
| cze | 12 | HP_and_AHP_Non_Physics_Authors | 36 |
| fin | 11 | HP_and_AHP_Non_Physics_Authors | 37 |
| isr | 11 | HP_and_AHP_Non_Physics_Authors | 37 |
| pol | 10 | HP_and_AHP_Non_Physics_Authors | 39 |
| vnm | 10 | HP_and_AHP_Non_Physics_Authors | 39 |
| nor | 9 | HP_and_AHP_Non_Physics_Authors | 41 |
| bgd | 9 | HP_and_AHP_Non_Physics_Authors | 41 |
| are | 9 | HP_and_AHP_Non_Physics_Authors | 41 |
| rou | 8 | HP_and_AHP_Non_Physics_Authors | 44 |
| mac | 6 | HP_and_AHP_Non_Physics_Authors | 45 |
| aut | 6 | HP_and_AHP_Non_Physics_Authors | 45 |
| phl | 6 | HP_and_AHP_Non_Physics_Authors | 45 |
| col | 5 | HP_and_AHP_Non_Physics_Authors | 48 |
| chl | 5 | HP_and_AHP_Non_Physics_Authors | 48 |
| qat | 5 | HP_and_AHP_Non_Physics_Authors | 48 |
| irl | 5 | HP_and_AHP_Non_Physics_Authors | 48 |
| nga | 4 | HP_and_AHP_Non_Physics_Authors | 52 |
| nzl | 4 | HP_and_AHP_Non_Physics_Authors | 52 |
| lbn | 4 | HP_and_AHP_Non_Physics_Authors | 52 |
| srb | 3 | HP_and_AHP_Non_Physics_Authors | 55 |
| isl | 3 | HP_and_AHP_Non_Physics_Authors | 55 |
| cyp | 3 | HP_and_AHP_Non_Physics_Authors | 55 |
| mar | 3 | HP_and_AHP_Non_Physics_Authors | 55 |
| bgr | 3 | HP_and_AHP_Non_Physics_Authors | 55 |
| jor | 3 | HP_and_AHP_Non_Physics_Authors | 55 |
| lux | 2 | HP_and_AHP_Non_Physics_Authors | 61 |
| omn | 2 | HP_and_AHP_Non_Physics_Authors | 61 |
| brn | 2 | HP_and_AHP_Non_Physics_Authors | 61 |
| svk | 2 | HP_and_AHP_Non_Physics_Authors | 61 |
| gha | 2 | HP_and_AHP_Non_Physics_Authors | 61 |
| ltu | 2 | HP_and_AHP_Non_Physics_Authors | 61 |
| tun | 2 | HP_and_AHP_Non_Physics_Authors | 61 |
| mex | 2 | HP_and_AHP_Non_Physics_Authors | 61 |
| kwt | 1 | HP_and_AHP_Non_Physics_Authors | 69 |
| geo | 1 | HP_and_AHP_Non_Physics_Authors | 69 |
| per | 1 | HP_and_AHP_Non_Physics_Authors | 69 |
| lva | 1 | HP_and_AHP_Non_Physics_Authors | 69 |
| dji | 1 | HP_and_AHP_Non_Physics_Authors | 69 |
| ukr | 1 | HP_and_AHP_Non_Physics_Authors | 69 |
| grd | 1 | HP_and_AHP_Non_Physics_Authors | 69 |
| hun | 1 | HP_and_AHP_Non_Physics_Authors | 69 |
| mus | 1 | HP_and_AHP_Non_Physics_Authors | 69 |
| syr | 1 | HP_and_AHP_Non_Physics_Authors | 69 |
| pse | 1 | HP_and_AHP_Non_Physics_Authors | 69 |
| ken | 1 | HP_and_AHP_Non_Physics_Authors | 69 |
| eth | 1 | HP_and_AHP_Non_Physics_Authors | 69 |
| ecu | 1 | HP_and_AHP_Non_Physics_Authors | 69 |
| uzb | 1 | HP_and_AHP_Non_Physics_Authors | 69 |
| kaz | 1 | HP_and_AHP_Non_Physics_Authors | 69 |

Supplementary Table 3. Number of non-Physics EP authors in each calendar year for the 8 countries with the highest fold-increases between 2016 and 2022.

| Country | Year | EP authors (non-Physics) |
| --- | --- | --- |
| tha | 2022 | 19 |
| sau | 2022 | 69 |
| rus | 2022 | 13 |
| pak | 2022 | 17 |
| kor | 2022 | 26 |
| ita | 2022 | 62 |
| ind | 2022 | 51 |
| esp | 2022 | 23 |
| tha | 2021 | 9 |
| sau | 2021 | 34 |
| rus | 2021 | 10 |
| pak | 2021 | 14 |
| kor | 2021 | 24 |
| ita | 2021 | 60 |
| ind | 2021 | 33 |
| esp | 2021 | 28 |
| tha | 2020 | 4 |
| sau | 2020 | 21 |
| rus | 2020 | 5 |
| pak | 2020 | 6 |
| kor | 2020 | 14 |
| ita | 2020 | 47 |
| ind | 2020 | 21 |
| esp | 2020 | 19 |
| tha | 2019 | 3 |
| sau | 2019 | 14 |
| rus | 2019 | 4 |
| pak | 2019 | 1 |
| kor | 2019 | 7 |
| ita | 2019 | 28 |
| ind | 2019 | 17 |
| esp | 2019 | 14 |
| tha | 2018 | 3 |
| sau | 2018 | 7 |
| rus | 2018 | 2 |
| pak | 2018 | 2 |
| kor | 2018 | 5 |
| ita | 2018 | 24 |
| ind | 2018 | 16 |
| esp | 2018 | 11 |
| tha | 2017 | 2 |
| sau | 2017 | 5 |
| rus | 2017 | 3 |
| pak | 2017 | 3 |
| kor | 2017 | 3 |
| ita | 2017 | 16 |
| ind | 2017 | 4 |
| esp | 2017 | 3 |
| tha | 2016 | 1 |
| sau | 2016 | 6 |
| rus | 2016 | 2 |
| pak | 2016 | 3 |
| kor | 2016 | 5 |
| ita | 2016 | 9 |
| ind | 2016 | 5 |
| esp | 2016 | 2 |
| tha | 2015 | 1 |
| sau | 2015 | 6 |
| rus | 2015 | 1 |
| pak | 2015 | 5 |
| kor | 2015 | 5 |
| ita | 2015 | 10 |
| ind | 2015 | 3 |
| esp | 2015 | 2 |
| tha | 2014 | 1 |
| sau | 2014 | 5 |
| rus | 2014 | 1 |
| pak | 2014 | 2 |
| kor | 2014 | 3 |
| ita | 2014 | 13 |
| ind | 2014 | 7 |
| esp | 2014 | 1 |
| sau | 2013 | 3 |
| pak | 2013 | 1 |
| kor | 2013 | 2 |
| ita | 2013 | 13 |
| ind | 2013 | 1 |
| esp | 2013 | 2 |
| tha | 2012 | 2 |
| sau | 2012 | 3 |
| pak | 2012 | 1 |
| kor | 2012 | 6 |
| ita | 2012 | 10 |
| ind | 2012 | 4 |
| esp | 2012 | 3 |
| sau | 2011 | 3 |
| rus | 2011 | 1 |
| pak | 2011 | 1 |
| kor | 2011 | 9 |
| ita | 2011 | 4 |
| ind | 2011 | 8 |
| esp | 2011 | 3 |
| rus | 2010 | 2 |
| pak | 2010 | 2 |
| kor | 2010 | 6 |
| ita | 2010 | 3 |
| ind | 2010 | 7 |
| rus | 2009 | 1 |
| pak | 2009 | 2 |
| kor | 2009 | 1 |
| ita | 2009 | 5 |
| ind | 2009 | 2 |
| esp | 2009 | 1 |
| tha | 2008 | 1 |
| kor | 2008 | 3 |
| ita | 2008 | 3 |
| ind | 2008 | 1 |
| esp | 2008 | 2 |
| tha | 2007 | 2 |
| ind | 2007 | 5 |
| esp | 2007 | 2 |
| tha | 2006 | 1 |
| ita | 2006 | 3 |
| ind | 2006 | 2 |
| esp | 2006 | 1 |
| esp | 2005 | 1 |
| sau | 2004 | 1 |
| rus | 2004 | 2 |
| ita | 2004 | 1 |
| esp | 2004 | 1 |
| ind | 2003 | 1 |
| rus | 2002 | 2 |
| ita | 2002 | 2 |
| ind | 2002 | 1 |
| rus | 2001 | 1 |
| ita | 2001 | 1 |

Supplementary Table 4. Number of HP, AHP, and EP authors in each scientific field

| Scientific Field | HP authors | AHP authors | EP authors |
| --- | --- | --- | --- |
| Physics & Astronomy | 10441 | 8588 | 12624 |
| Enabling & Strategic Technologies | 299 | 490 | 605 |
| Information & Communication Technologies | 229 | 292 | 396 |
| Chemistry | 155 | 248 | 307 |
| Engineering | 189 | 265 | 334 |
| Agriculture, Fisheries & Forestry | 106 | 93 | 143 |
| Mathematics & Statistics | 28 | 30 | 48 |
| Clinical Medicine | 523 | 929 | 1091 |
| Earth & Environmental Sciences | 43 | 57 | 79 |
| Built Environment & Design | 5 | 6 | 9 |
| General Science & Technology | 3 | 3 | 5 |
| Public Health & Health Services | 9 | 20 | 23 |
| Biomedical Research | 38 | 72 | 86 |
| Economics & Business | 9 | 13 | 20 |
| Biology | 20 | 21 | 38 |
| Psychology & Cognitive Sciences | 3 | 1 | 3 |
| Social Sciences | 2 | 3 | 4 |
| Philosophy & Theology | 0 | 0 | 0 |
| Historical Studies | 0 | 0 | 0 |
| Communication & Textual Studies | 0 | 0 | 0 |
| General Arts, Humanities & Social Sciences | 0 | 0 | 0 |
| Undetermined | 0 | 0 | 0 |
| Visual & Performing Arts | 0 | 0 | 0 |
